# PTRNAmark: an all-atomic distance-dependent knowledge-based potential for 3D RNA structure evaluation

**DOI:** 10.1101/076000

**Authors:** Yi Yang, Qi Gu, Ya-Zhou Shi

## Abstract

RNA molecules play vital biological roles, and understanding their structures gives us crucial insights into their biological functions. Model evaluation is a necessary step for better prediction and design of 3D RNA structures. Knowledge-based statistical potential has been proved to be a powerful approach for evaluating models of protein tertiary structures. In present, several knowledge-based potentials have also been proposed to assess models of RNA 3D structures. However, further amelioration is required to rank near-native structures and pick out the native structure from near-native structures, which is crucial in the prediction of RNA tertiary structures. In this work, we built a novel RNA knowledge-based potential— PTRNAmark, which not only combines nucleotides’ mutual and self energies but also fully considers the specificity of every RNA. The benchmarks on different testing data sets all show that PTRNAmark are more efficient than existing evaluation methods in recognizing native state from a pool of near-native states of RNAs as well as in ranking near-native states of RNA models.

---

In this work, we build an all-heavy-atom knowledge-based statistical potential called Precision Training RNA Mark (PTRNAmark). Unlike the aforementioned knowledge-based potentials that only consider non-bond interactions which belongmutual nucleotides, a new energy contribution based on the inside of nucleotide is involved in PTRNAmark. Furthermore, to consider the specificity of physical interaction, PTRNAmark is trained twice by two sorts of training sets, which is unchanging and changing respectively. For a decoy, firstly PTRNAmark is trained by a constant training set, like the aforementioned knowledge-based potentials, then PTRNAMark is trained by another training set which is some structures, originating from decoys, that are the lowest energy ranked by first time scoring. We think that this method could fully consider the characteristic of every RNA model and the specificity of physical interaction. It turns out that PTRNAmark performs better than 3DRNAScore, RASP, KB potentials and ROSETTA in ranking a tremendous amount of near-native RNA tertiary structures as well as recognizing native state from a pool of near-native states of RNAs.

## MATERIALS AND METHODS

The steps for generating PTRNAmark are as follows. First, we design the functional form of PTRNAmark from Boltzmann distribution, which merges nucleotides’ mutual and self-energies. Second, in order to train the parameters of the scoring function, which have been used to score for the decoys firstly, we select a training set of non-redundant RNA tertiary structures in which the structure features are representative and the structures of high similarity and low quality are removed. Third, we use the two test sets, which are occurring now, to test the accomplishment of PTRNAmark. Here, for every decoy, PTRNAmark is trained by some structures of the lowest energy which are from decoys and are scored by the PTRNAmark that is building in the second step. And, we use different metrics to compare the performance of PTRNAmark with other scoring methods. The more point of each building step of PTRNAmark is described in the follows.

## Generation of RNA potential

Our knowledge-based potential PTRNAMark is made of two terms. The first term based on the distance between any two non-bonded heavy atoms located at different nucleotides in the molecule, and the second term based on distance between any two non-bonded heavy atoms located at inside nucleotides in the molecule. According to three assumptions which were pointed out by Samudrala [27], the total energy score of a given RNA sequence S_q_ with conformation C_P_ is calculated by

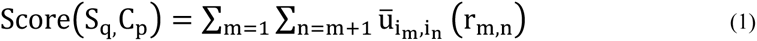

Where r_m,n_ are the distance between mth and nth atoms, and i_m_ and i_n_ are the residue-specific atom types, respectively.

## The energy term of mutual nucleotide

The knowledge-based potentials were derived based on the Boltzmann or Bayesian formulations. For the atomic distance-specific contact potentials, the potential can be written as:

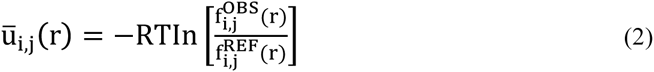

where R and T are Boltzmann constant and Kelvin temperature, respectively. 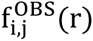 is the observed probability of atomic pairs (i, j) within a distance bin r to r+dr in experimental RNA conformations. 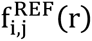 is the expected probability of atomic pairs (i, j) in the corresponding distance from random conformations without atomic interactions, which is so-called reference state. Here atomic pair (i, j) runs through all the atomic pairs in the RNA chain except for those pairs of the same nucleotides. Because of the reason that the equal size of datasets is used for calculating 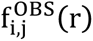 and 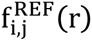 in RNA statistical potentials at present, the probabilities in Eq.(2) can be replaced by the frequency counts of atomic pairs:

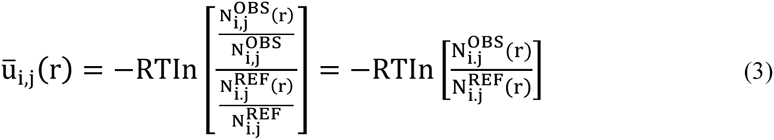

Here, 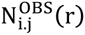 is the observed number of atom pairs (i, j) at the distance r in experimental RNA structures. 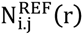 is the expected number of atomic pairs (i, j) if there were no interactions between atoms. 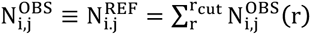 is the total number of atomic pairs (i, j) in thestructure samples, where r_cut_ is the cutoff distance. Cut off distance means the maximum value of the distance d in the process of statistics, and we find that when cut off distance is 20 Å, the number of atom pair observed is maxed, so we take 20 Å to be cut off distance in PTRNAmark. The counts of 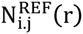 could not be compiled from experimental structures directly. It relies on which reference state we choose. The RASP and 3DRNAScore potentials both used averaged (RAPDF) reference states [27] and the KB potentials used quasi-chemical (KBP) [29] approximation reference states [30]. Our statistical potential chooses the averaged reference state, which ignores the type of atom. In averaged reference state,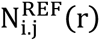 can be calculated as follows [27]:

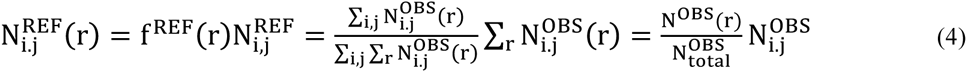

Here 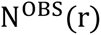 is the counts of observed contacts between all pairs of atom types at a particular distance r. 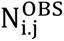 is the number of the occurrence of atom pairs of types i and j in whole distance region. 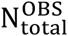 is the total number of contacts between all pairs of atom types summed over all distance r, it means the total counts. So the first term of functional form of PTRNAmark can be written as:

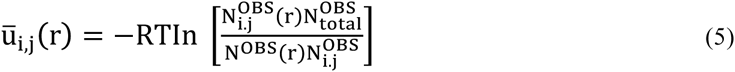

The energy term of nucleotide’s inside

We use not only mutual nucleotide potential, but also the inside potential of nucleotide, involving all atom pairs excluding these atom pairs which belong to bond stretching and angle bending in four RNA nucleotide inside. First, we calculated their statistical distribution over the training set. Once we get their statistical distributions, just like the mutual nucleotide potential, the potential can be written as:

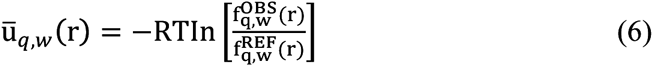

where R and T are Boltzmann constant and Kelvin temperature, respectively. 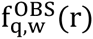 is the observed probability of atomic pairs (q, w) within a distance bin r to r+dr in experimental RNA conformations. 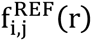 is the expected probability of atomic pairs (q, w) within a distance bin r to r+dr in a reference state. Here atomic pair (q, w) runs through all the atom-pairs precluding these atom pairs which belong to bond stretching and angle bending in four RNA nucleotide inside. Then we could also get the second term of functional form of PTRNAmark in the same way as the mutual nucleotide potential energy, it can be written:

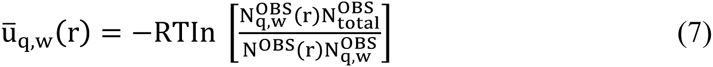

## Combination of the two energy terms

In PTRNAMark the two energy terms are combined together to get the final total energy:

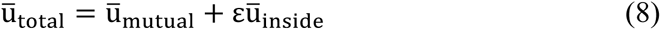

where 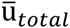 is the total energy, 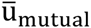 is the energy of mutual nucleotide 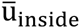 is the energy of nucleotide's inside, and ɛ is the weight. To get an appropriate value for ɛ, first we generate five decoy sets by using EFOLD program[31](PDBID:1MSY, 1ZIH, 1KKA, 1Q9A, 255D), then we calculated each decoy’s RMSD and the energy score, after that we maximize the ES by trying using a series of ɛ value. In the end, we find the optimized ɛ value is 0.15(see supplementary data VIII).

Our all-heavy-atom distance-dependent potential utilizes all the 85 atom types in the four nucleotides: 22 atom types in adenine (A), 20 atom types in cytosine (C), 23 atom types in guanine (G) and 20 atom types in uracil (U). In general, the distance-dependent statistical potential just considers nonbonding interactions between the different nucleotide. For PTRNAmark, the first term considers the nonbonding interaction between different nucleotide, and the second term includes all interactions of nucleotide inside except for bond stretching and angle bending interaction. so the atom pair in which the two atoms belong to the homogeneous nucleotide would not be considered in the first term and the second term only considers the atom pair which belongs to nucleotide’s inside except for bond stretching and angle bending.

For discrete statistics of data, the size of the bin has a great influence on the probability distribution. If the bin width is oversized, the result would be not truly precise. When the bin width is undersized, an unsuitable and artificial discontinuity of the result will occur, because of none or little samples located in certain bins. For protein potential, Sippl [26] used a bin width of 1 Å, Samudrala used a bin width of 1 Å and then carried out spline fitting [27]. For RNA potential, Capriotti (RASP) also used a bin width of 1 Å, Bernauer (KB) used a Dirichlet process mixture model, which leads to analytically differentiable potential functions, rather than fixed binning and spline fitting. Yi Xiao’s group (3DRNAScore), used a bin width of 0.3 Å, studying Scott’s work [32] in 1979 and extracting the bin width for 3dRNAscore from experimental structure. Here, considering that there is an observed diversity of the number of atom pairs in different bin width, we use the varying bin width. When the distance of atom pair < 3 Å, the bin width is 1 Å. If the distance of atom pair >= 3 Å, the bin width is 0.3 Å.

As for the problem of sparse data, in 1990, Sippl developed a method to address this problem. He approximated the genuine frequency by the sum of the total densities and the statistical frequencies [26]. Yi Xiao’s group (3DRNAScore) utilize a penalty to solve this problem, giving a penalty to the total energy score when the distance between two atoms is < 3 Å. In our PTRNAmark, we use a constant value to solve this problem.

## REFERENCE

1. Gesteland, R.F., Cech, T.R. & Atkins, J.F. The RNA World: The Nature of Modern RNA Suggests a Prebiotic RNA World (Cold Spring Harbor Laboratory Press, Cold Spring Harbor, New York, USA, 2006).

2. Guttman,M. and Rinn,J.L. (2012) Modular regulatory principles of large non-coding RNAs. Nature, 482, 339–346.

3. Dethoff,E.A., Chugh,J., Mustoe,A.M. and Al-Hashimi,H.M. (2012)Functional complexity and regulation through RNA dynamics.Nature, 482, 322–330.

4. Laing,C. and Schlick,T. (2011) Computational approaches to RNA structure prediction, analysis, and design. Curr. Opin. Struct. Biol., 21, 306–318.

5. Cao,S. and Chen,S.J. (2011) Physics-based de novo prediction of RNA 3D structures. J. Phys. Chem. B, 115, 4216–4226.

6. Das,R., Karanicolas,J. and Baker,D. (2010) Atomic accuracy in predicting and designing noncanonical RNA structure. Nat.Methods, 7, 291–294.

7. Frellsen,J., Moltke,I., Thiim,M., Mardia,K.V., Ferkinghoff-Borg,J. and Hamelryck,T. (2009) A probabilistic model of RNA conformational space. PLoS Comput. Biol., 5, e1000406.

8. Sharma,S., Ding,F. and Dokholyan,N.V. (2008) iFoldRNA: three-dimensional RNA structure prediction and folding. Bioinformatics, 24, 1951–1952.

9. Parisien,M. and Major,F. (2008) The MC-Fold and MC-Sym pipeline infers RNA structure from sequence data. Nature, 452, 51–55.

10. Martinez,H.M. Jr, Maizel,J.V. and Shapiro,B.A. (2008) RNA2D3D: a program for generating, viewing, and comparing 3-dimensional models of RNA. J. Biomol. Struct. Dyn., 25, 669–683.

11. Das,R. and Baker,D. (2007) Automated de novo prediction of native-like RNA tertiary structures. Proc. Natl. Acad. Sci. U.S.A.,104, 14664–14669.

12. Zhao,Y., Huang,Y., Gong,Z., Wang,Y., Man,J. and Xiao,Y. (2012) Automated and fast building of three-dimensional RNA structures. Sci. Rep., 2, 734.

13. Zhao,Y., Gong,Z. and Xiao,Y. (2011) Improvements of the hierarchical approach for predicting RNA tertiary structure. J. Biomol. Struct. Dyn., 28, 815–826.

14. Zhang,Y. (2008) Progress and challenges in protein structure prediction. Curr. Opin. Struct. Biol., 18, 342–348.

15. Zhou,H. and Skolnick,J. (2011) GOAP: a generalized orientation-dependent, all-atom statistical potential for protein structure prediction. Biophys. J., 101, 2043–2052.

16. Adhikari,A.N., Freed,K.F. and Sosnick,T.R. (2013) Simplified protein models: Predicting folding pathways and structure using amino acid sequences. Phys. Rev. Lett., 111, 28103.

17. Capriotti,E., Norambuena,T., Marti-Renom,M.A. and Melo,F. (2011) All-atom knowledge-based potential for RNA structure prediction and assessment. Bioinformatics, 27, 1086–1093.

18. Bernauer,J., Huang,X., Sim,A.Y.L. and Levitt,M. (2011) Fully differentiable coarse-grained and all-atom knowledge-based potentials for RNA structure evaluation. RNA, 17, 1066–1075.

19. Norambuena,T., Cares,J.F., Capriotti,E. and Melo,F. (2013) WebRASP: a server for computing energy scores to assess the accuracy and stability of RNA 3D structures. Bioinformatics, 29, 2649–2650.

20. Sim,A.Y.L., Schwander,O., Levitt,M and Bernauer,J. (2012) evaluating mixture models for building rna knowledge-based potentials. J. Bioinform. Comput. Biol., 10, 1241010.

21. Olechnovic,K. and Venclovas,C. (2014) The use of interatomic contact areas to quantify discrepancies between RNA 3D models and reference structures. Nucleic Acids Res., 42, 5407–5415.

22. Melo,F., Nchez,R. and Sali,A. (2002) Statistical potentials for fold assessment. Protein Sci., 11, 430–448.

23. Neal,R.M. (2000) Markov chain sampling methods for Dirichlet process mixture models. J. Comp. Graph. Stat., 9, 249–265.

24. J. Wang, Y. Zhao, C. Zhu, Y. Xiao, 3dRNAscore: a distance and torsion angle dependent evaluation function of 3D RNA structures, Nucleic acids research, 43 (2015) e63.

25. H. Deng, Y. Jia, Y. Wei, Y. Zhang, What is the best reference state for designing statistical atomic potentials in protein structure prediction, Proteins, 80 (2012) 2311–2322.

26. Sippl,M.J. (1990) Calculation of conformational ensembles from potentials of mena force: an approach to the knowledge-based prediction of local structures in globular proteins. J. Mol. Biol., 213, 859–883.

